# Lipid metabolic pathways determine phage infectivity in *Mycobacterium abscessus*

**DOI:** 10.64898/2026.03.25.714171

**Authors:** Mireia Bernabéu-Gimeno, Pilar Domingo-Calap

## Abstract

*Mycobacterium abscessus* is a rapidly growing non-tuberculous mycobacterium with high intrinsic antibiotic resistance, requiring innovative therapeutics. During infection, smooth and rough colony morphotypes can coexist in the human body, participating in the pathogenesis. Despite its increasing clinical importance and phage therapy being a last resort treatment recently, the genetic basis of phage interaction in *M. abscessus* remains poorly understood. Previous work has focused largely on rough strains or the surrogate host *Mycobacterium smegmatis*. Here, we isolated novel phages that efficiently infect both morphotypes, allowing us to characterize phage-resistant mutants from paired smooth and rough clinical isolates to determine the genetic basis of the infectivity. Integrating whole-genome sequencing, transcriptomics, and phage susceptibility and adsorption assays, we deeply analyzed 30 phage-resistant variants and found that resistance trajectories were shaped primarily by host morphotype rather than by the selecting phage. We confirmed the TPP locus as a conserved determinant of phage infection in both morphotypes and identified previously undescribed hotspot mutations in *furB* and *nrnA*, together with a multi-gene deletion, in smooth-derived resistant variants. Whereas the TPP locus and the multi-gene deletion represent stable genomic changes affecting lipid-associated loci, frameshift mutations in *furB* and *nrnA*, not previously linked to lipid metabolism, were accompanied by broad transcriptional rewiring of lipid-related genes. Resistance was mutation-dependent and consistently associated with impaired phage adsorption, indicating an early block in infection. Together, these findings show that phage recognition in *M. abscessus* is shaped by mycomembrane lipid architecture rather than a single dedicated receptor and uncover regulatory and metabolic pathways with implications for more durable phage-based therapies.

## Introduction

*Mycobacterium abscessus* is a rapidly growing non-tuberculous mycobacterium and an emerging multidrug-resistant pathogen that poses a major threat to individuals with underlying lung disease, particularly with cystic fibrosis^1,2^. Mycobacteria possess a distinctive mycolic-acid-rich outer membrane that underpins their intrinsic resistance to antibiotics^3^. Despite the rising global prevalence of *M. abscessus,* its molecular biology remains comparatively understudied relative to other mycobacteria, notably *M. tuberculosis*^1^. A defining feature of *M. abscessus* biology is its ability to adopt two colony morphotypes, smooth and rough, which differ primarily in the presence of surface-exposed glycopeptidolipids (GPLs)^4–6^. Smooth variants dominate during early colonization, whereas rough variants frequently arise during chronic infection through mutations in GPL biosynthesis or transport genes^7–9^. Both morphotypes often coexist within a single patient, reflecting ongoing within-host evolution^10^. In addition, the molecular determinants that govern its interaction with bacteriophages (or phages) remain poorly defined. Notably, colony morphotype is the strongest known predictor of phage susceptibility, with rough variants generally being more permissive to infection, whereas smooth isolates display marked resistance and inefficient killing^11^.

Nearly all characterized and most therapeutic phages were isolated in surrogate hosts (mostly *M. smegmatis*) and belong to the cluster AB, including Muddy as a reference phage^12^. Despite the clinical success of phage therapy against *M. abscessus*^13,14^, mechanistic insights into infection remain largely undescribed, with all investigations confined to *M. smegmatis* or rough *M. abscessus* strains^15–20^. Host factors such as the nucleoid-associated protein Lsr2 modulate infection dynamics, with phages preferentially adsorbing at sites of new cell-wall synthesis^16^. In addition, trehalose polyphosphate (TPP) has been proposed as a co-receptor for Muddy, with its gp24 minor tail protein described as a key determinant of host specificity^15^. However, the identity of bona fide receptors is still undetermined. Critically, the genetic basis of phage infection in smooth clinical isolates, the morphotype that predominates during early human infection^4,6^, remains largely unexplored, despite recent evidence that smooth-to-rough morphotype switching, mostly driven by GPL-associated mutations, can mediate resistance to a lysogenic phage^21^.

Here, we uncover previously unrecognized mutational hotspots that impair phage adsorption by characterizing phage-resistant mutants in both smooth and rough *M. abscessus* variants obtained from the same patient. For that purpose, we isolated and characterized novel phages capable of efficiently infecting both morphotypes. These previously undescribed determinants map predominantly to genes involved in lipid metabolic pathways distinct from canonical GPL biosynthesis, supporting a model in which the composition of *M. abscessus* cell envelope plays a key role in phage entry.

## Results

### Three novel AB mycophages providing new receptor-binding protein diversity

To identify phages capable of infecting clinical rough and smooth *M. abscessus* isolates, we performed phage hunting using *M. smegmatis* as a surrogate host. Three novel mycobacteriophages, Falla, Goset and Gegant, were isolated. Transmission electron microscopy revealed siphovirus morphology, characterized by icosahedral capsids (∼100–150 nm) and long, non-contractile tails (Fig. 1A). Whole-genome sequencing showed genome sizes with Falla having a slightly shorter genome (47,046 bp) compared to Goset and Gegant (47,519 bp and 47,687 bp). Comparative BLASTn analysis against the Actinobacteriophages database (February 2026) assigned all three phages to the cluster AB, displaying high nucleotide similarity to previously described members, including the reference phage Muddy (GenBank NC_022054.2). Goset and Gegant showed 99% coverage with 95.76% and 95.64% nucleotide identity, respectively, and Falla showed 97% coverage with 95.57% identity. Pharokka annotation of the phages together with other related phages (FF47, 8UZL, Maco6 and Muddy), followed by pham-based clustering, placed Falla, Goset and Gegant within the strictly lytic AB cluster, with pairwise proteomic equivalence quotient (peq) >0.9 across all compared phages (Fig. 1B). Manual curation confirmed the absence of temperate markers and enabled detailed, gene-by-gene comparisons among our phages and with Muddy (Fig. 1C). Falla differed from Goset/Gegant by missing five predicted CDSs of unknown function and by encoding a unique 96-aa protein without functional annotation. This protein had no pham association within the cluster AB, although BLASTp against the Actinobacteriophage Proteins Database showed limited similarity to other unknown proteins from different clusters. Complete phage annotation, including correspondence between local phams and Actinobacteriophage database phams (date: 06 March 2026), assigned using Pham2Phage is detailed in Table S1. Currently described as the main receptor-binding protein in other AB phages, the minor tail protein gp24 was analyzed in depth, revealing high variability among Falla, Goset and Gegant, with substitutions mainly toward the 3′ end. Specifically, relative to Falla gp24, the other two shared four substitutions (E616A, K638E, Q654E and T657S), with Goset having five additional substitutions (I320V, E441Q, A442D, A531S and V554I) and Gegant eight (A502D, T504D, I516M, N610S, D659N, T661A, V714A and T722I). Homology to the mycophage Douge central fiber protein gp20 (GMQE=0.33; QSQE=0.19; identity=23.88) enabled prediction of the gp24 homotrimer with AlphaFold3 (ipTM=0.55, pTM=0.57, Fig. 1D). Variable residues accumulated mainly in the loop regions that structurally aligned to the C-terminal domain of Douge gp20, except for three substitutions in β-sheets (positions 610, 638 and 657) and one substitution in an α-helix (position 320).

**Fig. 1.**
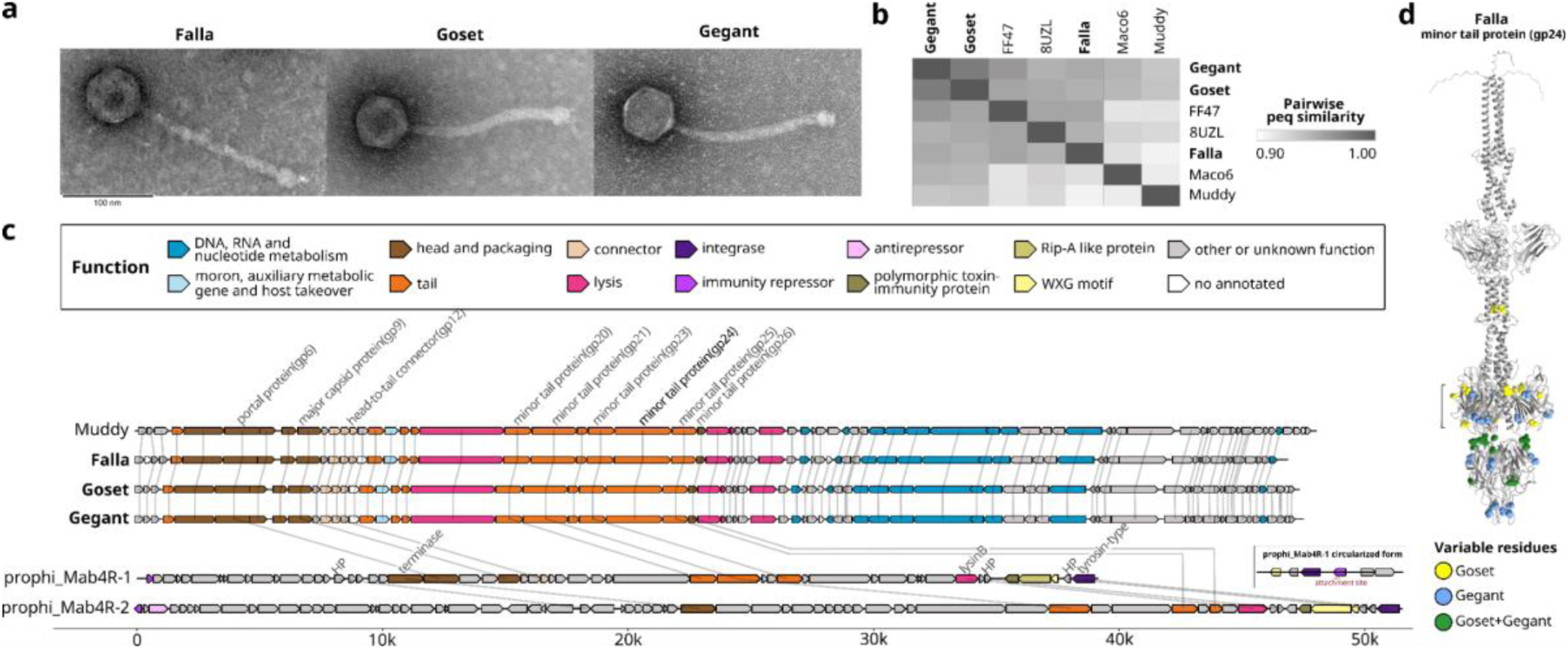
Characterization of the novel mycophages Falla, Goset, and Gegant. **a. Transmission electron microscope images.** Scale bars are indicated. b. Phage cluster assignment. Proteome-based clustering, including representative cluster AB phages (FF47, 8UZL, Maco6, and Muddy). Colour based on pairwise proteomic equivalence quotient (peq) similarity. c. Phage genomic comparison. Manually accurate annotation of Falla, Goset, and Gegant and pham-related genes with the reference phage Muddy and Mab4R/Mab5S prophages (prophi_Mab4R-1 and prophi_Mab4R-2) is shown. The circular variant of prophi_Mab4R-1 (encoding an immunity repressor) is alongside its chromosomally integrated form. d. Structural analysis of minor tail protein gp24. Phage Falla gp24 homotrimer model predicted with AlphaFold3 and visualized with ChimeraX is displayed, including amino acid substitutions distinguishing Goset and Gegant from Falla. The region aligning to the C-terminal domain of gp20 is marked.

### Phage infectivity is preserved across smooth and rough morphotypes

Smooth and rough morphotypes are commonly found intrapatient. Under this scenario, Mab4R and Mab5S were coexisting variants isolated from the same patient, differing in morphotype (Fig. 2A). Under the conventional classification, Mab4R is “rough” (dry colonies with irregular margins), whereas Mab5S is “smooth” (shiny colonies despite irregular edges). To directly compare phage susceptibility of these morphotype-differing variants, we evaluated the three newly isolated phages in semi-solid and liquid infection assays. Spot tests revealed comparable lytic activity of phages Falla, Goset, and Gegant against the clinical smooth-rough pair (Fig. 2A), indicating that morphotype did not restrict plaque formation. In liquid-killing assays, the three phages effectively reduced bacterial loads ranging from 5 × 10² to 5 × 10⁷ CFU/mL of both variants in all tested phage concentrations (5 to 5 × 10⁸ PFU/mL) (Fig. 2B). Whole-genome sequencing followed by pangenome-based phylogeny confirmed the close genetic relationship between Mab4R and Mab5S and their proximity to *M. abscessus subsp. abscessus* ATCC19977 (Fig. 2C). Specifically, both Mab4R and Mab5S chromosomes encode the macrolide-resistance gene *erm (41)*, five complete defense systems, and two prophages (prophi_Mab4R-1 and prophi_Mab4R-2) (Fig. 2D). Moreover, each strain harbors a 22,349 bp plasmid (defined as Mab4R plasmid) encoding a MobF-family relaxase. Compared to Mab4R, Mab5S holds nine non-synonymous mutations (including the GPL-locus gene *lys2B*/*MAB_4098c*), three additional changes in non-coding regions and a 1,980-bp insertion carrying three genes encoding a nucleoside-diphosphate-sugar epimerase (*MAB_4711*), an AraC-family protein (*MAB_4712*) and a Zn-dependent glyoxalase (*MAB_4713*) (Fig. 2D). Given that prophages are thought to be major contributors to mycobacterial defence, prophi_Mab4R-1 and prophi_Mab4R-2 were included in the comparative phage analysis (Fig. 1C). Interestingly, minor tail proteins from prophages pham-clustered with AB lytic phages: gp20 and gp23 with prophi_Mab4R-1, gp25 and gp26 with prophi_Mab4R-2 and gp23 with both prophages. Moreover, several head and packaging proteins, including the portal protein (gp6), the major capsid protein (gp9), and the head-to-tail connector (gp12) from the AB lytic phages, shared the assigned pham with prophi_Mab4R-1. In Mab4R, prophi_Mab4R-1 was detected circular with a ∼30-fold coverage increase, consistent with spontaneous excision (Table S3). Comparison between inserted and viral forms of prophi_Mab4R-1 suggested the presence of an integration-dependent immunity system^22^. In contrast, no induction was detected in Mab5S, which carries a single non-coding change located 5 bp upstream of the prophi_Mab4R-1 immunity repressor gene. Complete information about the Mab4R and Mab5S comparison is provided in Table S2.

**Fig. 2.**
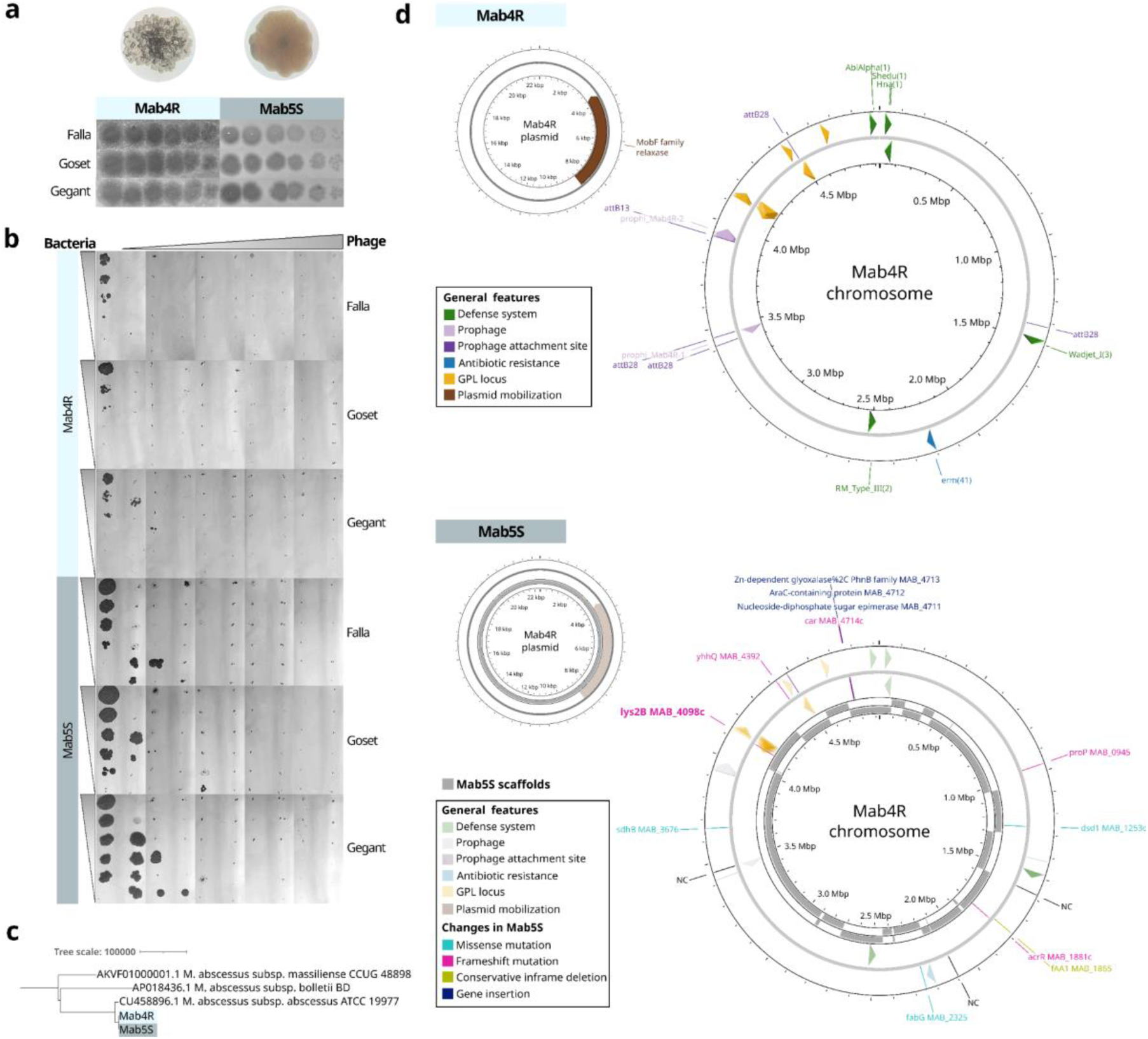
Phage-susceptibility in rough and smooth *M. abscessus* variants. **a. Bacterial morphotype and spot test.** Colony morphology in LB + CaCl₂ plates is shown above the spot assay with 2 µL of serial dilutions of phages Falla, Goset, and Gegant (10^10^ PFU/mL) spotted onto Mab4R (rough) and Mab5S (smooth) lawns. **b. Liquid killing assay.** 100 µL of Mab4R or Mab5S (10–10⁶ CFU) was mixed with 100 µL of LB + 5 mM CaCl₂ (no-phage control) or 100 µL of phage Falla, Goset, or Gegant (10¹–10⁸ PFU) and incubated for 48 h at 37 °C with shaking. The resulting growth after spotting 2 µL of each mixture onto LB + CaCl₂ plates and incubating for 7 days is shown. **c. Taxonomic classification.** Phylogeny was inferred with Gubbins using a Panaroo core-genome alignment of Mab4R, Mab5S, and representative strains from each *M. abscessus* subspecies. **d. Genomic analysis.** The Mab4R assembly with hybrid reads is shown with selected features highlighted. For Mab5S, scaffolds from the SKESA assembly were mapped to the Mab4R and shown in grey. Non-synonymous mutations and changes in non-coding regions are indicated. The gene most likely associated with the morphotype switch is highlighted in bold.

### Mutational hotspots underlying phage resistance across morphotypes

To better understand phage infectivity in smooth and rough colonies, a methodology was designed to isolate stable phage-resistant variants. Incubation of 10⁸ CFU of each morphotype was challenged with the phages independently at a multiplicity of infection (MOI) of 10 during different time periods (2, 5 and 7 d), followed by multiple serial washes and single-colony purification. This allowed the selection of free-phage-resistant variants from the original phage-selected population, smooth and rough (Figure 3A). Specifically, 10 colonies were screened per incubation time (2, 5 or 7 days), bacterium (Mab4R or Mab5S) and phage (Falla, Goset or Gegant). Mab4R yielded 18 phage-free variants with increased resistance (7 selected with Falla, 7 with Goset and 4 with Gegant), while in Mab5S, only one resistant colony was initially phage-free. After additional passages to remove the remaining phage, 9 additional Mab5S-resistant colonies were obtained: 2 from Falla, 2 from Goset and 7 from Gegant. These 7 Gegant-derived variants comprised 4 colonies from the initial phage-selected population and 3 α-β pairs (two variants derived from the same original colony but selected in additional passages owing to distinct morphotypes and/or phenotypes). No visible morphotype changes were detected in any Mab4R variants. In contrast, half of the smooth phage-resistant variants exhibited consistent, perceptible changes in colony morphology.

**Fig. 3.**
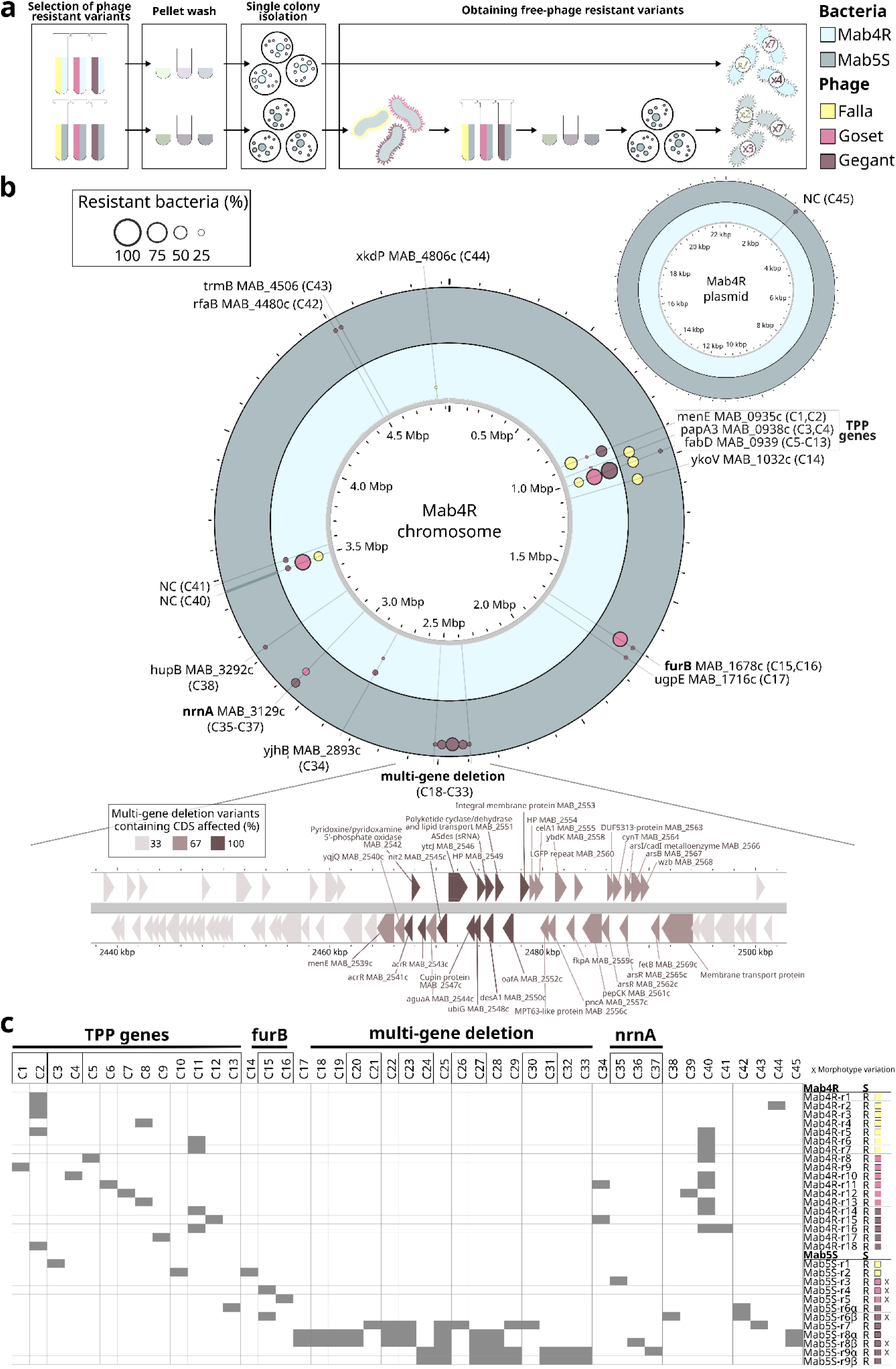
Rough and smooth *M. abscessus* phage-resistant variants. **a. Simplified experimental design**. Mab4R (rough) or Mab5S (smooth) was incubated with a 10-fold higher titer of phage Falla, Goset, or Gegant at 37 °C with shaking. Single colonies from the resulting populations were isolated, and those with reduced susceptibility and no residual phage were selected. For Mab5S, additional growth–wash single-colony purification cycles were needed. **b. Mutational hotspots of phage-resistant variants.** For both Mab4R- and Mab5S-resistant variants selected with each phage, the genomic distribution of the observed changes and the frequency of single colonies affected are shown. For the multi-gene deletion, due to the variable length of the chromosomal deletion, additional frequency analysis is included. **c. Matrix of genomic alterations in phage-resistant variants**. Changes affecting the same gene are shown together with those affecting the major hotspots (TPP genes, *furB*, *nrnA*, or the multi-gene deletion) marked in bold.

Sequencing analysis of the 30 phage-resistant variants, defined as ‘Mab4R-rX’ and ‘Mab5S-rX’, was performed to evaluate the molecular determinants of the phage resistance (Fig. 3B). All Mab4R variants carried non-synonymous mutations in TPP biosynthesis genes, menE (*MAB_0935c*), papA3 (*MAB_0938c*) and fabD (*MAB_0939*), suggesting a mutational hotspot in this region, with the highest diversity in fabD (nine distinct changes: C5–C13). While three Mab5S-resistant variants also carried non-synonymous mutations in the TPP biosynthesis genes *MAB_0938c* or *MAB_0939*, as observed in the rough variants, interestingly, three additional hotspots were noticed. These mutations affected the *furB* (*MAB_1678c*), the *nrnA* (*MAB_3129c*) genes, and a variable-length chromosomal deletion (25,863–39,437 bp) located between the coordinates 2,439,523 and 2,501,235. This deletion consistently removed nine genes (*MAB_2542* and *MAB_2546*-*MAB_2553*), the *MAB_2550c*-associated ASdes sRNA, and fully or partially deleted three additional genes (*MAB_2541c*, *MAB_2543c* and *MAB_2545c*). Moreover, variant-specific extensions included other genes located upstream or downstream (*MAB_2508*-*MAB_2539c*, *MAB_2544c* and/or *MAB_2554*-*MAB_2578c*). Across both Mab4R- and Mab5S-resistant variants, additional non-synonymous mutations affected seven chromosomal genes, together with a plasmid mutation in a non-coding region. A complete description is provided in Table S4, with variant-specific supporting data in Table S5. However, all these mutations always co-occurred with at least one major hotspot mutation (TPP genes, *furB*, *nrnA* or the multi-gene deletion) (Fig. 3C). Despite the detection of two non-coding changes affecting the prophi_Mab4R-1 in 9 Mab4R-resistant variants (C40 and C41), none of the prophages appeared to be contributing to the resistance in any of the 30 phage-resistant variants.

### Phenotypic colony changes link phage resistance and morphotype transition

As previously mentioned, half of the smooth phage-resistant variants showed colony morphology variation, which might be contributing to phage resistance. To further evaluate the phenotypic effect of each mutational hotspot, spot and reverse-spot assays across phages at different titres ranging from 10⁴ to 10⁸ PFU were performed, revealing a mutation-dependent resistance (Fig. 4A). Mab5S-resistant variants containing the affecting-*furB* changes (Mab5S-r4 and Mab5S-r5 with no additional mutations) initially showed the strongest resistance phenotype (full phenotype), with no plaque formation and sustained growth against 10⁶ PFU of all three phages in the reverse-spot test. However, this phenotype was unstable, and the variants exhibited another phenotype in which resistance was subtle and almost indistinguishable from the wild type (partial phenotype). Interestingly, all *furB* frameshift variants showed colonies with increased edge irregularity compared to Mab5S. This change in morphotype also occurred in variants containing affecting-*nrnA* changes, whose colony had a rounder shape with less irregular edges. These variants showed partial resistance, visible as slightly increased plaque turbidity in spot tests and, more consistently, as reduced growth restriction compared with the wild type in reverse-spot assays as phage titres increased. Multi-gene deletion variants showed a clear resistant phenotype by reverse-spot assays, most evident at the 10⁶ PFU exposure, where Mab5S-r8α retained visible growth throughout the dilution series. However, no change was observed in the standard spot test. Interestingly, for the *furB*, *nrnA* and multi-gene deletion variants, resistance levels were comparable across Falla, Goset and Gegant. In contrast, TPP-pathway mutants showed stronger resistance to Falla than to Goset/Gegant, regardless of the phage used for selection, consistent with the reduced titre of Falla in these variants as observed in the spot tests. Differences in morphotype-associated susceptibility were clear in the reverse-spot test assay of TPP-pathway variants. In rough mutants, resistant growth relative to the wild type was detectable only at 10⁴ PFU for Goset or Gegant. At higher phage exposure, bacterial growth was observed only with Falla, and the increased exposure also revealed differences among the affected TPP genes, with *MAB_0938c* mutants showing slightly greater resistance than *MAB_0935c* mutants. In the smooth variants, differences in susceptibility between the wild-type and resistant variants became apparent only in the 10⁶ and 10⁸ PFU spot tests, which also revealed greater resistance to Goset than to Gegant. Moreover, the three α-β pairs, consisting of two variants derived from the same original colony selected in the additional passages, further support the described mutation-dependent phenotype (Fig. 4B). In all the cases, both variants shared core mutations but also included changes consistent with their distinctive morphotypes and phage resistance. Specifically, Mab5S-r6α and Mab5S-r6β shared the change C42 but differed in additional *furB*- and TPP-pathway-specific mutations correlating with the colony morphology and resistant phenotype. Moreover, Mab5S-r8β and Mab5S-r9α, which displayed a rounder colony appearance compared to their pair counterparts (Mab5S-r8α and Mab5S-r9β, respectively), carried an *nrnA*-affecting mutation. No differences in resistance in these two α-β pairs were appreciated, possibly due to the higher degree of resistance conferred by the common multi-gene deletion.

**Fig. 4.**
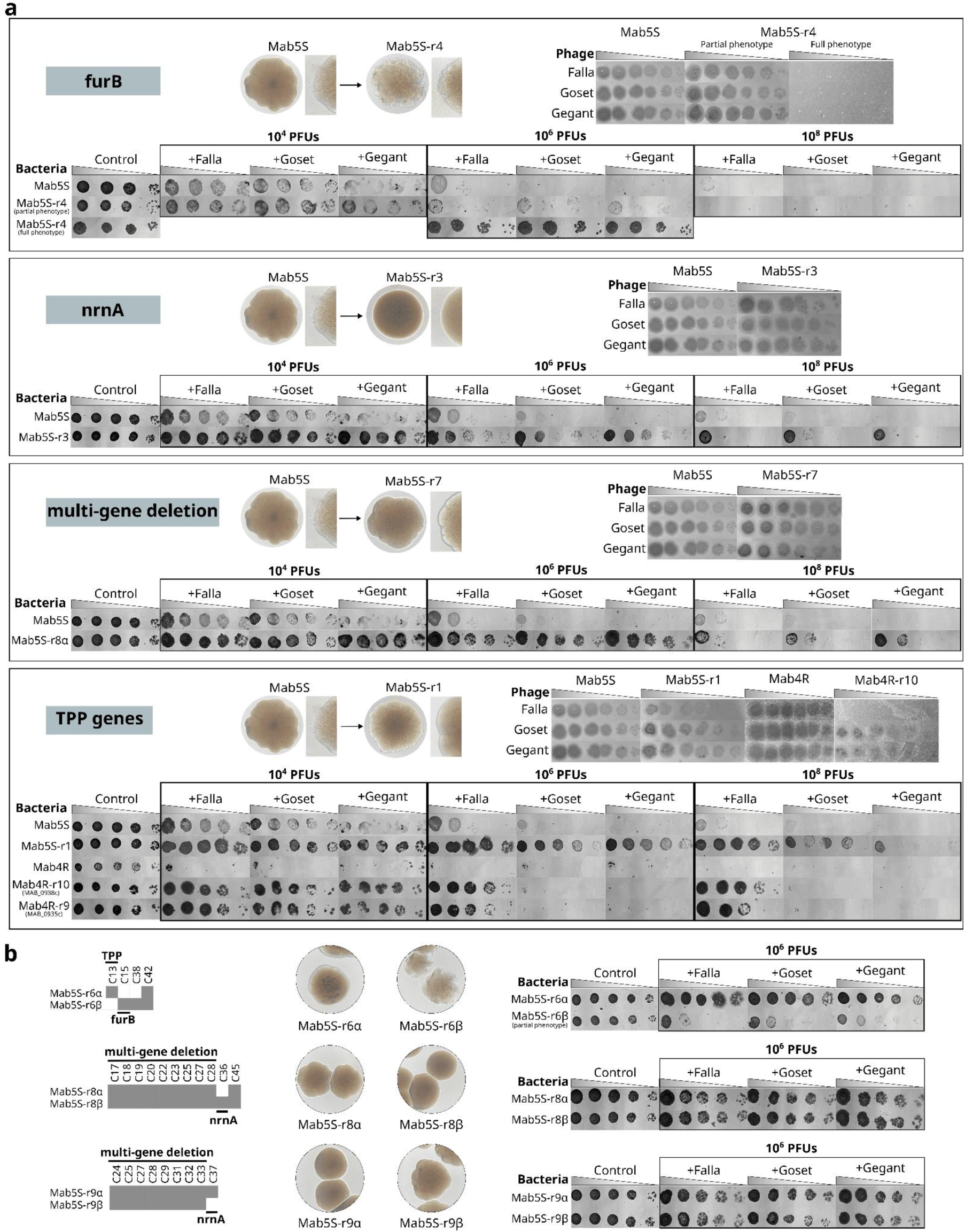
Mutation-specific phenotypes of phage-resistant variants. **a. Phage susceptibility and colony morphotype.** For *furB*, *nrnA*, the multi-gene deletion and TPP genes, a representative variant is shown, including colony appearance on LB + CaCl₂ and classical spot and reverse-spot assays performed across increasing phage titres (10⁴–10⁸ PFU). The corresponding wild-type strain used for selection is shown for comparison. For *furB*, two phage-associated phenotypes (full and partial phenotypes) are included. **b. Comparison of colony-derived α-β pairs.** For each pair, consisting of two variants derived from the same original colony selected in the additional passages, differences in genomic alternations, colony morphotype and/or phage resistance by reverse-spot test are displayed.

### Mutations associated with impaired phage adsorption and lipid metabolic remodelling

To determine whether resistance impaired phage recognition, adsorption assays were conducted using Mab5S and its derived resistant variants. This strain was selected due to the broader mutational diversity identified in the smooth background, including variants analogous to those observed in rough isolates. Under conditions optimized for adsorption to Mab5S, specific binding of Falla, against which TPP-pathway mutants displayed the strongest resistance, was evaluated with the parental Mab5S strain serving as a control for comparison. Adsorption efficiency was quantified as the ratio of free phage titres after 1 h relative to time 0 (T1/T0), where lower values indicate greater phage adsorption. Collectively, the resistant variants showed significantly impaired adsorption compared with the wild type (0.59 ± 0.33 versus 0.085 ± 0.03; Welch’s t-test, t = 7.29, df = 25.1, p = 1.2 × 10⁻⁷) (Fig. 5A), including variant-to-variant differences consistent with the observed spot test resistance, with multi-gene deletion and TPP mutants displaying stronger adsorption defects than *furB* or *nrnA* mutants. In three of the five multi-gene deletion variants, this adsorption defect was statistically significant (Mab5S-r8A p = 0.0052; Mab5S-r8B p = 0.0087; Mab5S-r9A p = 0.0228) based on Holm-adjusted pairwise contrasts versus Mab5S following Welch’s ANOVA. The *nrnA* mutant Mab5S-r3 showed a milder adsorption defect (0.31 ± 0.02), and as observed in the susceptibility assays, no adsorption differences were appreciated between the members of α-β pairs differing in the additional *nrnA*-affecting mutation (Mab5S-r8α and Mab5S-r8β; Mab5S-r9α and Mab5S-r9β). Despite phenotypic variability, the *furB* mutants consistently showed an adsorption defect, similar in magnitude to *nrnA* (0.26 ± 0.09), even when exhibiting the lowest resistance phenotype (partial phenotype).

**Fig. 5.**
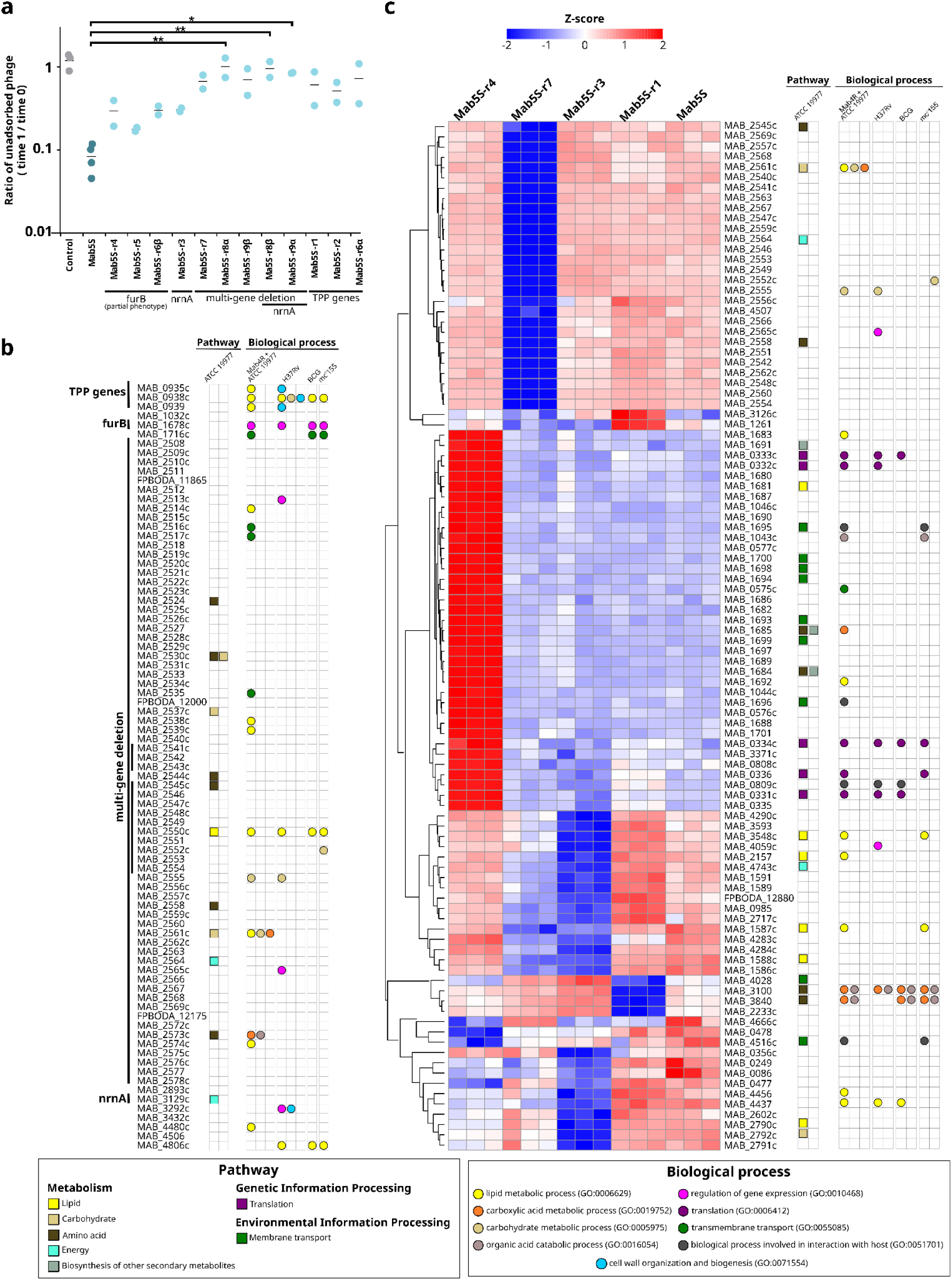
Resistance association with impaired phage adsorption and bacterial lipid-related pathways. **a. Phage Falla adsorption to *M. abscessus* variants.** Ratios (free phage titre at 1 h relative to time 0) were computed per biological replicate from three technical replicates by averaging log₁₀ (PFU/mL) and back-transforming the difference. The y-axis is shown on a logarithmic scale, and variants are grouped by mutational hotspot (*furB*, *nrnA*, multi-gene deletion, and TPP genes). Control corresponds to the phage incubated in parallel in the absence of bacteria. Statistical significance versus Mab5S (smooth wild-type as a control) is indicated (* = P < 0.05; ** = P < 0.01). **b. Functional annotation of bacterial mutated genes.** Curated ATCC 19977 pathway genes from KEGG were transferred to equivalent Mab4R/Mab5S proteins (≥70% query coverage and ≥90% identity). Mab4R/Mab5S GO biological-process Bakta annotations were expanded by transferring GO terms from the UniProt equivalent ATCC 19977 protein. Additional functional information was obtained via OMA orthology between ATCC 19977 and *M. tuberculosis* H37Rv, *M. bovis* BCG Pasteur 1173P2, and *M. smegmatis* mc²155 to retrieve UniProt GO terms. Using all GO terms, aggregated GO groups were generated with R (AnnotationDbi) and assigned to each protein. **c. RNA-seq evidence of lipid-related transcriptional changes.** Genes meeting an expression threshold (logCPM ≥ 1) and significant in at least one comparison (FDR < 0.05) were ranked by the maximum absolute log₂ fold-change across contrasts. The top 100 were plotted using log2-CPM expression values scaled per gene (row Z-scores). Functional annotations for genes included in the heatmap were obtained as described for the mutated genes in Fig. 5B.

Functional analysis using Gene Ontology terms and KEGG pathways linked the mutational hotspots to lipid metabolism (Fig. 5B). Specifically, all the TPP genes (*MAB_0935c*, *MAB_0938c* and *MAB_0939*) are associated with lipid-related processes, and the common deleted region includes *MAB_2550c*, encoding desA1 enzyme involved in lipid metabolism in *M. abscessus* and orthologs. Moreover, the variant-specific deletion extensions always included at least two additional genes potentially involved in lipid metabolism (*MAB_2514c*, *MAB_2538c* and *MAB_2539c*; *MAB_2539c* and *MAB_2561c*; or *MAB_2561c* and *MAB_2574c*). Interestingly, *MAB_2539c* and *MAB_2574c* both encode an o-succinybenzoic acid–CoA lyase MenE. For variants without direct mutations in lipid-metabolism genes (*furB* and *nrnA*), RNA sequencing suggested differential expression of lipid-related genes (Fig. 5C). Specifically, comparing with Mab5S, the frameshift mutation in the *furB* transcriptional repressor (carried by Mab5S-r4) was associated with overexpression of a 37-gene set, including three lipid-related genes (*MAB_1683*, *MAB_1681* and *MAB_1692*), and the *nrnA*-affecting variant Mab5S-r3 showed a decreased expression of many lipid-related genes (*MAB_3548c*, *MAB_2157*, *MAB_4456*, *MAB_4437*, *MAB_2790c*, *MAB_1587c* and *MAB_1588c*). Notably, these last two genes were also underexpressed in the multi-gene deletion variant Mab5S-r7, in which no decrease in *MAB_2550c* expression was observed, unlike for the rest of the deleted genes. Other hotspot co-occurring affected genes, such as *MAB_4480c* and *MAB_4806c*, are likewise involved in lipid metabolism, as well as in cell-wall organization and biogenesis (*MAB_3292c*), a biological process that is also associated with the TPP genes. Secondarily, the affected genes also appeared to participate in additional biological processes, including: carbohydrate metabolism (*MAB_0938c*, *MAB_2530c*, *MAB_2537c*, *MAB_2552c*, *MAB_2555*, *MAB_2561c* and *MAB_2792c*), membrane transport (*MAB_2516c*-*MAB_2517c*, *MAB_2535*, *MAB_1716c*, *MAB_4028*, *MAB_1693*-*MAB_1696*, *MAB_1698*-*MAB_1700* and *MAB_4516c*), regulation of gene expression (*MAB_2513c*, *MAB_2565c*, *MAB_3292c* and *MAB_4059c*) and amino acid metabolism and translation (*MAB_2524*, *MAB_2530*, *MAB_2544*-*MAB_2545*, *MAB_2558*, *MAB_2573c*, *MAB_0331c*-*MAB_0334c*, *MAB_0336*, *MAB_1684*-*MAB_1685*, *MAB_3100* and *MAB_3840*). Collectively, these findings identify lipid-associated genetic alterations as key determinants of phage adsorption and resistance in *M. abscessus*.

## Discussion

The ability of phages to target multidrug-resistant *M. abscessus* remains a critical but poorly understood aspect of phage-based therapeutic development. Our study confirmed that phage infectivity is not restricted by colony morphotype and identified lipid-associated genetic alterations as key determinants of resistance and adsorption efficiency. The identification of previously undescribed mutational hotspots linked to phage resistance provides a new genetic framework for understanding mycophage-host interactions. In particular, the association between resistance phenotypes, adsorption impairment, and lipid metabolic remodelling supports a model in which cell envelope composition plays a central role in phage entry dynamics. With previous studies suggesting the involvement of surface-associated molecules in mycophage infection, our results extend this concept by revealing specific genetic loci potentially underlying these processes.

Our experimental approach was based on isolates harboring a mobilizable plasmid and prophages, situating resistance evolution within a clinically represented genomic context^22–24^. By selecting phage-resistant variants in both morphotypes, we uncovered distinct adaptive trajectories determined primarily by host background. In the rough isolate, resistance converged on the TPP locus, consistent with prior evidence implicating TPP as a co-receptor for phage infection^15^. In contrast, smooth-derived variants revealed a broader mutational landscape, including a multi-gene deletion and non-synonymous substitutions in *furB* and *nrnA*, genes not previously linked to mycophage susceptibility. These findings suggest that, in addition to the TPP-related pathways defined in rough strains, alternative lipid-associated or regulatory mechanisms can modulate phage adsorption in smooth backgrounds. Morphotype instability in clinical isolates^25^ likely contributes to this adaptive plasticity. The emergence of *nrnA* mutants with altered colony appearance illustrates how resistance and morphotype transitions may be intertwined, rather than independent processes, as described recently for smooth phage-resistant variants switching to rough morphotypes^21^. Although additional hotspots were not detected in rough-derived variants, their absence could reflect differences in selective dynamics rather than strict morphotype restriction. The reduced efficiency of phage clearance in smooth cultures may prolong selective pressure, facilitating exploration of a wider mutational landscape. Interestingly, resistance conferred by *furB*, *nrnA*, and the multi-gene deletion was broadly similar across the three phages, while the TPP mutants were more resistant to Falla than to Goset or Gegant. A plausible explanation for these phage-specific differences is variation in the receptor-binding protein gp24, since a few mutations in this region enable Muddy to overcome TPP-associated defense^15^. Indeed, our phages expand the known diversity of the gp24. Variable residues in this protein were accumulated in a region structurally similar to the C-terminal domain of the central fiber protein gp20, a receptor-binding protein reported to bind fragments of major mycobacterial surface antigens (LAM/AM and α-glucan)^26^, supporting a role in specificity and receptor engagement. However, targeted gp24 mutagenesis might be done to confirm this hypothesis and evaluate the role of each specific residue.

Evaluation of phage susceptibility in mycobacteria remains technically challenging, as bacterial aggregation can introduce variability and limit reproducibility across experimental platforms^27^. In this context, we found reverse-spot tests across graded phage concentrations particularly informative, as increased selective pressure revealed subtle phenotypic differences that were not apparent in standard spot tests. In all resistant variants, the phenotype correlated with the underlying genetic alteration, with impaired phage adsorption emerging as the shared functional consequence. The degree of adsorption defect closely paralleled the degree of the resistance phenotype, indicating that resistance in our system is primarily mediated at the earliest stage of infection. The most pronounced phenotypes were associated with stable genomic changes affecting lipid-associated loci, including genes involved in TPP biosynthesis and a multi-gene deletion encompassing the desaturase *desA1* (*MAB_2550c*). In *M. smegmatis* and *M. tuberculosis, desA1* is essential and participates in mycolic-acid biosynthesis^28^, and its associated ASdes sRNA has been proposed to regulate fatty acid desaturase desA2^29^. The apparent tolerance of this deletion in *M. abscessus* variants is consistent with previous suggestions that *desA1* may not be strictly essential in this species^30^. However, because the deletion extended to at least two additional lipid-related genes, including *MAB_2539c* and *MAB_2574c*, both encoding an O-succinybenzoic acid–CoA lyase MenE, the individual contribution of each locus remains to be elucidated.

On the other hand, although *furB* (encoding a transcriptional repressor) and *nrnA* (nanoRNase/pAp phosphatase) are not classically associated with lipid metabolism, frameshift mutations in these genes were linked to significant transcriptional variation of diverse lipid-associated genes in our exploratory RNAseq analysis. Notably, the *furB* frameshift conferred the most pronounced resistance phenotype, characterized by the complete absence of plaque formation. As the *furB* frameshift resulted in derepression of a 37-gene regulon, several of which are involved in lipid metabolism, we hypothesize that this transcriptional rewiring alters the composition, architecture, or accessibility of critical mycomembrane components required for phage adsorption, potentially including the phage receptor itself. Such mechanisms would represent a regulatory route to phage resistance that is mechanistically distinct from canonical receptor loss. Alternatively, given *FurB*’s conserved role in zinc homeostasis in other mycobacterium species^31^, resistance may arise from altered metal availability, potentially triggering a previously unrecognized metal-dependent defense state. Although metal sequestration strategies are well described in host–*Mycobacterium* interactions^32^, their involvement in phage resistance has not been explored yet.

Taken together, these results reinforce the central role of cell envelope architecture in mediating phage adsorption in *M. abscessus.* Mycolic acids and associated lipid components are major determinants of membrane organization and surface accessibility, and even subtle perturbations in lipid composition could alter receptor exposure or membrane biophysical properties. In this context, the recurrent involvement of lipid-associated genes across independent resistant variants supports a model in which phage recognition is highly sensitive to the structural state of the mycomembrane rather than to a single dedicated receptor molecule. Although the multi-gene deletion precludes assignment of resistance to a single locus, the convergence on lipid metabolic pathways across distinct genetic events argues against a stochastic effect. Instead, these data suggest that remodelling of envelope-associated lipids represents a common adaptive route to phage resistance. Future targeted complementation and lipidomic analyses will be required to resolve the relative contribution of individual genes and to define the precise biochemical changes underlying impaired adsorption. On the whole, our findings confirm TPP as a conserved determinant for phage infection in both rough and smooth *M. abscessus* morphotypes and expand the landscape of bacterial determinants to include novel regulatory and metabolic pathways not traditionally linked to phage-host interaction. The identification of *furB*, *nrnA,* and genes encompassed within the multi-gene deletion highlights lipid-associated cell envelope modulation as a central axis of phage resistance. Importantly, these findings suggest that transcriptional regulators, often overlooked in phage-resistance screenings, may indirectly but profoundly influence phage susceptibility through secondary effects on membrane composition and cellular physiology.

From a translational perspective, these results have implications for phage therapy against *M. abscessus*. Regulatory or metabolic routes to resistance may be reversible, condition-dependent, or associated with fitness trade-offs, opening opportunities for combination strategies that exploit envelope vulnerabilities or metal homeostasis pathways. A deeper mechanistic understanding of the unique structural and regulatory features of the *M. abscessus* cell envelope will therefore be essential to rationally design durable phage-based interventions against this highly antibiotic-resistant pathogen.

## Materials and methods

### Bacterial strains and culture conditions

*M. smegmatis* mc^2^155 was obtained from the Spanish Type Culture Collection (CECT), and the isogenic clinical *M. abscessus* pair from the same patient (rough Mab4R, smooth Mab5S) was kindly provided by the Ramón y Cajal University Hospital (Madrid, Spain). Cultures were grown at 37 °C with shaking in Luria-Bertani broth supplemented with 5 mM CaCl_2_ (LB + CaCl_2_) for 48-72 h (*M. smegmatis*) or 5-7 d (*M. abscessus*). Morphotype was assigned by colony appearance on both LB + CaCl_2_ and blood agar plates.

### Phage isolation, amplification, and concentration

Phages were isolated from sewage water in Valencia (Spain) using stationary-phase *M. smegmatis*. Samples were centrifuged (3,900 x g, 10 min) and filtered (0.45 then 0.22 µm). Phages were recovered using the soft agar overlay method as previously described^33^ and purified by a triple plaque-to-plaque transfer. For amplification, phages were propagated in LB + CaCl₂ (48 - 72 h), lysates centrifuged (3,900 rpm, 5 min) and filtered (0.45 then 0.22 µm), and stocks stored at -70 °C. For characterization, filtered lysates were high-speed centrifuged (80,000 rpm, 2 h), and pellets resuspended in 200 µL SM buffer (50 mM Tris-HCl, pH 7.5; 8 mM MgSO_4_; 100 mM NaCl; 0.01% gelatin, wt/vol). Transmission electron micrographs were obtained from purified phage lysates at the Prince Felipe Research Centre (CIPF).

### Bacterial susceptibility assays

Susceptibility of bacterial isolates to phages was assessed by spot tests (2 µL of serial phage dilutions starting at 10^10^ PFU/mL on bacterial lawns; 5-7 d growth at 37 °C), reverse-spot tests (2 µL serial stationary-phase bacteria, 5 × 10⁷ - 5 × 10⁶ CFU/mL, on phage lawns containing 10⁴, 10⁶, or 10⁸ PFU; 5-7 d growth at 37 °C), and liquid killing assays as described^14^ with minor modifications: 100 µL bacteria (10 - 10^6^ CFU) with 100 µL LB + CaCl₂ or 100 µL phage (1 - 10^8^ PFU), incubated for 48 h at 37 °C with shaking; then 2 µL were spotted onto LB + CaCl₂ plates, incubated for 7 d, and photographed. Assays were performed with at least two biological replicates.

### Isolation of phage-resistant variants

*In vitro* selection used 10⁸ CFU of wild-type clinical strains incubated at 37 °C with shaking for 2, 5, or 7 d with or without 10⁹ PFU of phages. Cultures were pelleted (3,900 x g, 10 min), resuspended in 1 mL, washed three times (14,000 x g, 5 min) in fresh LB + CaCl₂, streaked, and single colonies isolated; phage susceptibility was evaluated with resistance defined as reduced susceptibility compared to wild-type (lower phage titer, increased plaque turbidity, and/or increased survival in reverse-spot tests). Phage presence in resistant cultures was assessed by *M. smegmatis* spot tests and phage-specific PCR on supernatants. When detected, additional growth, pellet washes, and single-colony purification on LB + CaCl₂ were performed, followed by the susceptibility evaluation. Phage-free resistant isolates were defined as ’Mab4R-rX’ and ’Mab5S-rX’ and stored in 50% glycerol at -70 °C.

### Phage-adsorption assays

Conditions were optimized for ∼90% adsorption in Mab5S: 500 µL of 10x concentrated bacteria at OD_600_ = 0.25 (5·10⁸ CFU/mL) mixed with 50 µL phage at 10⁷ PFU/mL and incubated for 1 h at 37 °C with shaking. Before and after phage incubation, samples were centrifuged (14,000 x g, 1 min, 4 °C), supernatants serially diluted, and free phage titers quantified by plating on *M. smegmatis*. Adsorption was expressed as the log_10_ difference in phage titers (before/after incubation). A phage-only control (no bacteria) assessed titre loss due to phage instability. Each strain was tested in at least two biological replicates, each with three technical replicates; technical means were used as biological values. Data were plotted with ggplot2^34^ and analyzed in R^35^ using Welch’s ANOVA (Levene’s test p < 0.001), followed by Holm-adjusted contrasts comparing each variant to Mab5S.

### DNA extraction and sequencing

Phage DNA was extracted from concentrated stocks after host DNA removal as described before^33^. After capsid digestion with proteinase K, DNA was purified with DNA Clean & Concentrator-5 (Zymo Research) or the Maxwell PureFood GMO and Authentication kit (Promega). For *M. abscessus* isolates, pellets from 1 mL stationary-phase culture were resuspended in 270 µL 1x PBS, and DNA was extracted with the Maxwell RSC Pathogen Total Nucleic kit, including the lysozyme treatment. Libraries (Nextera XT) were sequenced on the Illumina MiSeq (2 x 250 bp phages; 2 x 300 bp bacteria). Mab4R was additionally sequenced on a MinION Mk1C (FLO-MIN114) using Rapid Barcoding Kit 24 v14 (SQK-RBK114-24), including the basecalling (fast model).

### Phage genomic characterization

Phage reads were quality-filtered with fastp^36^ (min_length: 50; trim_qual_right: 30; trim_qual_type: mean; trim_qual_window: 10) and assembled with SPAdes^37^ (--only-assembler). Preliminary cluster assignment was done based on the closest relatives identified by BLASTn^38^ against the Actinobacteria database^39^. Genomes were reoriented to the first Muddy gene and polished with Pilon^40^. Structural and functional annotation was performed with Pharokka^41^, including the preliminary cluster genomes. Local phamilies (“phams”) were defined with Phammseqs^42^ (--pangenome), and preliminary cluster assignment was checked with Phamclust^43^. Pharokka calls were manually curated to minimize differences between phages due to tool-derived artefacts. Annotated genomes are available in GenBank. Prophages were annotated with Depht^44^ and included in the Phammseqs analysis to assess shared evolutionary origin. Correspondence between local phams and Actinobacteria database phams was inferred with the custom pipeline Phage2Pham, which includes visualization with gggenomes^45^. Because Muddy gp24 is predicted to be the main receptor-binding protein, amino acid sequences were compared across phages: templates were identified with SWISS-MODEL^46^, and gp24 homotrimer was predicted with AlphaFold3^47^ (database date: 03 February 2026). Structures were inspected in ChimeraX^48^ (variable residues highlighted), and domain alignments were performed with Matchmaker tool (single chains).

### Bacterial genomic characterization

Quality filtering was performed with fastp^36^ (min_length: 70; cut_right_mean_quality: 20; cut_right_window_size: 3; detect_adapter_for_pe) for the Illumina reads and NanoFilt^49^ (-l 500 --headcrop 75 -q 10) for the Nanopore reads. Mab4R was hybrid-assembled with Unicycler^50^, and Illumina reads not mapping to the assembled chromosome were assembled with SPAdes^37^ to recover the plasmid. Reads unmapped to both the chromosome and plasmid were assembled with Unicycler to recover the circular prophiMab4R-1 genome. Chromosome and plasmid were annotated with Bakta^51^; resistance genes and defense systems were predicted using the web versions of ResFinder^52^ and DefenseFinder^53^, and prophages were screened with Depht^44^. Mab5S Illumina reads were assembled with SPAdes^37^ and SKESA^54^ in parallel, and both assemblies were used throughout; scaffolds were characterized as for Mab4R. The prophiMab4R-1 insertion site in Mab4R was inferred using Mab5S scaffolds and confirmed by manual insertion followed by read mapping and visualization. Low-complexity regions were identified with DustMasker^38^. A pangenome analysis of Mab4R and Mab5S used Panaroo^55^ core alignment (--clean-mode strict) followed by Gubbins^56^ recombination masking, using Bakta-annotated Mab4R, Mab5S (SKESA scaffolds) and representative *M. abscessus* subspecies genomes (GCF_000069185.1, GCF_000270665.1, GCF_003609715.1). The resulting tree was visualized with iTOL^57^. Orthology to the reference *M. abscessus subsp. abscessus* ATCC 19977 proteome (NC_010397.1) was inferred by BLASTp^38^, defining equivalents as >=70% query coverage and >=90% identity. Bakta GO^58^ biological-process annotations were expanded by transferring GO terms from UniProt database^59^ when an equivalent ATCC 19977 protein was found. Additional functional information was obtained by OMA orthology^60^ between ATCC 19977 and *M. tuberculosis* H37Rv, *M. bovis* BCG Pasteur 1173P2, and *M. smegmatis* mc2155 to retrieve UniProt GO terms (UP000001584, UP000001472 and UP000000757, respectively). Annotations were integrated in R (AnnotationDbi^61^) to reconcile orthology and generate aggregated GO groups; curated ATCC 19977 pathway genes were downloaded from KEGG^62^ and transferred to equivalent Mab4R/Mab5S proteins.

Wild-type strains and derived resistant variants were analyzed following a similar pipeline, described in Fig. S1. Read-based calling mapped filtered Illumina reads to the comparison genome with BWA^63^ to quantify coverage, called SNPs/small indels with Snippy^64^, GATK^65^, and Octopus^66^, and assembled unmapped reads with SPAdes^37^. Variants supported by at least two callers and outside DustMasker regions were retained. Assembly-based calling mapped SKESA and SPAdes assemblies to the comparison genome with minimap2^67^ and identified variants with svim-asm^68^; only calls present in both assemblies and outside DustMasker regions were kept. A deletion in five resistant isolates was validated by read mapping and PCR targeting the common deleted region (F: 5’-TGTGTCTCAGCGGTATTGGT-3’; R: 5’-TGCCCCTAATCAAGCGTCA-3’). One-base indels in MAB_1678c (three variants) were confirmed by PCR (F: 5’-TTCTGTTGATCGCCAAGGGG-3’; R: 5’-CTCATCAGCCACCTACG-3’) and Sanger sequencing. All variants were visualized in Proksee^69^ using the Mab4R genome as scaffold, and resistance-associated changes were also represented in a 0-1 matrix with Morpheus^70^.

### Bacterial RNA sequencing

Each bacterial strain was grown to OD_600_ = 0.7, and three technical replicates were used for RNA extraction using the Maxwell RSC miRNA Tissue kit (Promega), including the lysozyme treatment. Libraries were prepared with the RiboZero Plus rRNA depletion kit and sequenced on an Illumina NextSeq 500 (1 x 75 bp). Reads were mapped with Bowtie2^71^ to the Mab4R genome and the Mab5S-specific insertion; counts were generated with featureCounts^72^. Differential expression used edgeR^35,73^, treating each genotype (Mab5S and each variant) as a separate experimental condition: low expression genes were filtered (filterByExpr; grouping=genotype), library sizes normalized by the TMM method (calcNormFactors), a Mab5S reference matrix designed, dispersions estimated (estimateDisp), models fitted (glmQLFit) and quasi-likelihood F-tests per variant (glmQLFTest) on the relevant coefficient. For heatmaps, contrasts were combined, and genes with the minimum expression threshold (logCPM >= 1) and significant in at least one comparison (FDR < 0.05) were ranked by the maximum absolute log₂ fold-change across contrasts. The top 100 were plotted using log2-CPM expression values scaled per gene (row Z-scores).

## Notes

### Competing Interest Statement

P.D.-C. is cofounder and scientific advisor at Evolving Therapeutics.

### Funding

- Ministerio de Ciencia, Innovación y Universidades: CNS2023-145519, RYC2019-028015-I

